# Staging Alzheimer’s disease in the brain and retina of *B6.APP/PS1* mice by transcriptional profiling

**DOI:** 10.1101/741421

**Authors:** Sumana R. Chintapaludi, Asli Uyar, Harriet M. Jackson, Casey J. Acklin, Xulong Wang, Michael Sasner, Gregory W. Carter, Gareth R. Howell

## Abstract

Alzheimer’s disease (AD) is a common form of dementia characterized by amyloid plaque deposition, TAU pathology, neuroinflammation and neurodegeneration. Mouse models recapitulate some key features of AD. For instance, the B6.*APP/PS1* model (carrying human transgenes for mutant forms of *APP* and *PSEN1*) shows plaque deposition and associated neuroinflammatory responses involving both astrocytes and microglia beginning around 6 months of age. However, in our colony, TAU pathology, significant neurodegeneration and cognitive decline are not apparent in this model even at older ages. Therefore, this model is ideal for studying neuroinflammatory responses to amyloid deposition. Here, RNA sequencing of brain and retinal tissue, generalized linear modeling (GLM), functional annotation followed by validation by immunofluorescence (IF) was performed in B6.*APP/PS1* mice to determine the earliest molecular changes prior to and around the onset of plaque deposition (2-6 months of age). Multiple pathways were shown to be activated in response to amyloid deposition including the JAK/STAT and NALFD pathways. Putative, cell-specific targets of STAT3, a central component of the JAK/STAT pathway, were identified that we propose provide more precise options for assessing the potential for targeting activation of the JAK/STAT pathway as a treatment for human AD. In the retina, GLM predicted activation of vascular-related pathways. However, many of the gene expression changes comparing B6 with B6.*APP/PS1* retina samples occurred prior to plaque onset (2 months of age). This suggests retinal changes in B6.*APP/PS1* mice may be an artefact of overexpression of mutant forms of APP and PSEN1 providing limited translatability to human AD. Therefore, caution should be taken when using this mouse model to assess the potential of using the eye as a window to the brain for AD.

## INTRODUCTION

Alzheimer’s disease (AD) is a debilitating neurological disorder affecting as many as 10% of Americans over the age of 65 and 32% of people 85 or older [1]. The characteristic progressive decline in memory, learning, executive and visual functions is preceded by amyloid deposition, neurofibrillary tangles, neuroinflammation and the loss of cortical neurons and synapses in the brain [1]. Age is the strongest risk factor for AD. Individuals with familial AD carry high-penetrant mutations in APP or APP processing genes throughout their life and yet plaque deposition does not occur immediately, normally decades later. The same is true for sporadic or late-onset AD, which likely depends on the interaction of multiple genes, age, and environmental factors. Clinical AD progression is considered to occur in a standard sequence over years or decades: amyloid accumulation; tau-mediated neuronal synaptic and neuronal loss; changes in brain structure; and cognitive deficits [2]. Some mouse models relevant to AD also show an age-dependent disease onset and progression. For instance, the B6.*APP/PS1* model that carries mutations in both APP (*APPswe*) and the APP processing gene presenilin 1 (*PSEN1de9*), do not show plaque accumulations until they are about 6 months [3].

While AD-related pathology in the brain is well documented, the disease has also been reported to affect the retina, a developmental outgrowth of the brain [4]. Multiple human and animal studies have illustrated a correlation of AD neuropathology in the retina and the brain, including Aβ and tau accumulation, neuronal cell loss, and retinal ganglia cell (RGC) apoptosis [5]. Studies suggest that changes occur earlier in the retina than in other parts of the brain [5]. However, some studies suggest the value of using the retina to track AD pathology may be limited [6]. Therefore, the eye potentially represents a unique tissue that may serve as a valuable model for studying cerebral pathology and for discovering new biomarkers for AD. Given that AD pathology precedes the onset of the mental decline, the discovery of an early pathological endophenotype could be key for monitoring initial stages of the disease with the potential for uncovering pathophysiological mechanisms that trigger cognitive decline and dementia.

Understanding the triggering events of age-related diseases, such as AD, are challenging because of the lack of access to postmortem tissues at middle age and pre-symptomatic stages, and the asynchronous nature of disease onset in humans. Therefore, animal models are essential to study the early stages of disease initiation and early progression [7]. Gene expression profiling provides an unbiased approach for investigating the pathogenesis of complex diseases like AD. RNA-seq technology has the advantage of screening the expression of the whole genome making it one of the best tools for searching for novel biomarkers. Previously, we employed gene profiling to identify pathways impacted by plaque deposition in B6.*APP/PS1* mice [8]. Here we now explore the potential of applying sensitive computational approaches to directly stage brain and retinal changes during the onset and early stages of disease. These findings could be used to effectively translate the knowledge of these pathophysiological mechanisms of the disease into clinical applications.

## MATERIALS AND METHODS

### Mouse strains, breeding and husbandry

All procedures were approved by the Institutional Animal Care and Use Committee (IACUC) at The Jackson Laboratory and followed the Association for Research in Vision and Ophthalmology (ARVO) Statements for the Use of Animals in Ophthalmic and Vision Research. B6.Cg-Tg(APPswe, PSEN1dE9)85Dbo/Mmjax mice (hereafter called “B6.*APP/PS1”; MMRRC #34832-JAX*) [9] were obtained from The Jackson Laboratory and maintained in 14/10-hour light/dark cycle. *APP/PS1* and wild-type (WT) cohorts were generated by mating B6.*APP/PS1* hemizygous mice with C57BL/6J (B6; JAX #664) mice. Female mice between 2–6 months of age used in this study were housed under the same conditions in the same animal facility, minimizing environmental differences. All mice were maintained on Labdiet®5K52.

### Cohort generation and tissue harvesting

A total of 60 female mice were selected for transcriptional profiling of the brain: 10 at 2 months (5 WT, 5 B6.*APP/PS1*), 14 at 4 months (9 WT, 5 B6.*APP/PS1*), 24 at 5 months (13 WT, 11 B6.*APP/PS1*) and 12 at 6 months (7 WT, 5 B6.*APP/PS1*). Retinas were obtained from 22 mice: 10 at 2 months (5 WT, 5 B6.*APP/PS1*) and 12 at 6 months (7 WT, 5 B6.*APP/PS1*). Data and associated metadata have been submitted to Synapse (https://www.synapse.org/#!Synapse:syn18879982). At the above ages, left retinas and left brain hemispheres were dissected and snap frozen for preservation of RNA for gene expression studies. The right brain hemispheres and eyes were fixed in 4% paraformaldehyde overnight at 4°C and stored in 1× Phosphate-buffered solution (PBS) at 4°C for immunohistological studies.

### RNA Sequencing

Total RNA was extracted using Trizol (Invitrogen, CA). mRNA was purified from total RNA using biotin-tagged poly dT oligonucleotides and streptavidin-coated magnetic beads and quality was assessed using an Agilent Technologies 2100 Bioanalyzer. 2 × 100 bp paired end sequencing of RNA libraries were done on Illumina HiSeq 2000 at an average sequencing depth of ∼70 million reads per sample (**Fig S1**). Quality control of raw sequencing data was performed using FastQC tool. Reads were quality trimmed and filtered using Trimmomatic tool [10]. Reads passing the quality filtering were mapped to mm10 reference genome using ‘bowtie’ algorithm [11]. RNA-Seq by Expectation-Maximization (RSEM) software package was used to estimate expression levels for all genes in Transcripts Per Million (TPM) unit [12]. Principal Component Analyses (PCA) was used to identify clusters of samples based on transcriptomic patterns, and to detect potential outliers in the study cohort.

### Generalized linear model (GLM)

Genes were retained that met the following two criteria: TPM greater than 0 in at least 10% of samples; and maximum expression of the gene greater than 0.25 quantile of the complete dataset. Downstream analyses were conducted on the 10,843 genes that met these criteria. A generalized linear model (GLM) was used to identify genes with differential expression as a function of age, genotype, and age-genotype interaction. The samples were collected in two different experiments, generating a potential batch effect, and consequentially sample collection dates were included in the linear model. GLM was implemented in R Studio Version 1.1.456. We used FDR corrected p-value (qval) and r^2^ from the linear model to determine how well each gene’s expression across the samples could be fit by the predictors of the model, which were age, genotype and age-genotype interaction. For each gene, the GLM returned an estimate for each predictor’s effect size and corresponding significance. We used a relatively permissive threshold (qval < 0.25 & r^2^ > 0.3) to identify genes potentially fit by the model, and then categorized the significant genes further based on effects of individual predictors. Taking the WT as reference genotype, and 2 months as reference age, we extracted significant genes (p < 0.05) for transgene (Group APP), age (Age 4 months, Age 5 months and Age 6 months) and age-genotype interaction (Age 4 months: Group APP, Age 5 months: Group APP, Age 6 months: Group APP).

### Functional Enrichment Analysis

Functional enrichment analysis was performed using findGO.pl script within HOMER software [13]. A separate list of significant genes associated with each predictor was provided and the script was run with organism set to ‘mouse’. Enriched KEGG pathways from HOMER’s output were extracted and presented.

### Analysis with Cytoscape Software

The iRegulon plugin [14] in Cytoscape 3.2.0 was used to identify transcription factors and their targets predicted to be regulated by *STAT3* in mouse brain. iRegulon detects the TFs and co-regulated genes by scanning known TF-binding promoter motifs as well as the predicted motifs discovered from the Encyclopedia of DNA Elements (ENCODE) chromatin immunoprecipitation-sequencing data. All the settings were set as default.

### Immunofluorescence

Fixed brain and eyes were equilibrated in 15% sucrose at 4 °C overnight, then incubated in 30% sucrose. The tissues were then embedded in OCT (Sakura Finetek Europe B.V., Netherlands). Cryosections (12 μm sections for retinas and 30 μm for brain) were collected onto Superfrost Plus microscope slides (Fisher Scientific, Atlanta, GA). For immunostaining involving antibodies for vascular associated proteins in brain, sections were pretreated with Pepsin as previously described [91], others were left untreated. Sections were hydrated with H_2_O for 3 min at 37°C followed by treatment of the tissue with 0.5 mg/ml of Pepsin (Sigma-Aldrich) for 15 min at 37°C. Sections were then rinsed twice with 1X PBS at room temperature (RT) for 10 min. After Pepsin pretreatment, sections were rinsed once in 1X PBT (PBS + 1% Triton 100X) and incubated in primary antibodies diluted with 1X PBT + 10% normal goat or normal donkey serum over night at 4°C. Subsequently, sections were rinsed three times with 1X PBT for 10 min and incubated for two hours in the corresponding secondary antibodies (1:700, Invitrogen). Tissue was then washed three times with 1X PBT for 10–15 min, nuclei were stained with DAPI and mounted in Poly-aquamount (Polysciences). The following primary antibodies were used: chicken polyclonal anti-GFAP (1:250, Acris Antibodies); rabbit polyclonal anti-IBA1 (1:250, Wako); goat polyclonal anti-CD31 (1:40, R&D Technologies) and rabbit polyclonal anti-STAT3 (1:100, Cell Signaling Technologies), rabbit polyclonal anti-pSTAT3 (1:100, Cell Signaling Technologies), mouse monoclonal anti-β-Amyloid, 1-16 (1:1000, Biolegend). Sections were viewed, and images were obtained using the Leica SP5 confocal microscope. All microscope settings, including laser levels and gain, were held constant to allow for relative comparisons of signal intensity within and between experiments. Post image processing was performed in Fiji (http://fiji.sc/Fiji).

## RESULTS

### *Development of amyloid plaques in brain and retina in* B6.*APP/PS1 mice*

Our aim was to identify and compare expression changes in the brain and retina during the development of amyloid pathology in B6.*APP/PS1* mice. Previously, we reported plaque burden in B6.*APP/PS1* mouse brains using ThioS staining which stains for both neuritic plaques and tangles [8]. In this study, we used 6E10 staining which was specific to amyloid. In agreement with previous studies, B6.*APP/PS1* mice showed little evidence of plaques at 2 months of age but developed plaques by 6 months (Fig. 1). Aβ deposits were also observed in retinas from 2-6 months old B6.*APP/PS1* mice (Fig. 1). However, retinal Aβ deposits appeared morphologically different to their cerebral counterparts, comparatively smaller in sizes, diffuse and immature and primarily associated with retinal vasculature (Fig. 1).

**Figure 1.**
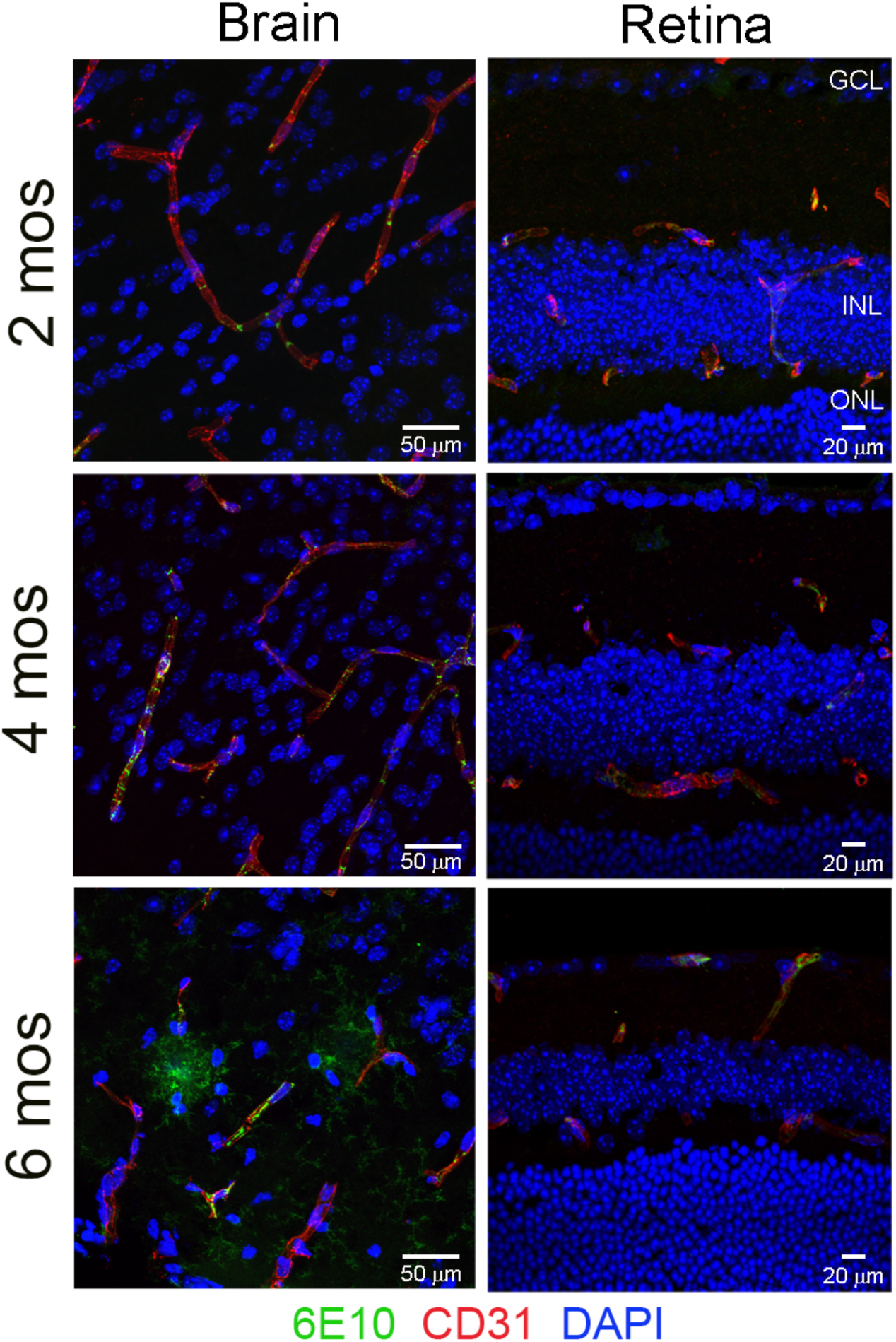
Amyloid plaques in female B6.*APP/PS1* mouse brain and retina between 2-6 months of age. Representative micrographs from paired brain and retina cross sections of B6.*APP/PS1* mice (n=4-6 per age group) at varying ages. In brain, little Aβ staining was observed at 2 and 4 mos (6E10; *green*), very rarely in endothelial cells (CD31; *red*). Aβ plaques positive for 6E10 staining (*green*) were commonly observed in the brain at 6 months of age. Scale bar: 50 μm. Retinal Aβ deposits (6E10) showed co-localization with CD31. Retinal Aβ deposits are apparent inside blood vessels, perivascular, and along the vessel walls. In contrast to the brain, little to no plaques were observed outside of vessels in the retina. TO-PRO-3 iodide staining labeled nuclei of all cells (*blue*). Abbreviations: GCL, ganglion cell layer; INL, inner nuclear layer; ONL, outer nuclear layer. Scale bar: 20 μm.

### Generalized linear modeling of brain transcriptomes from B6.APP/PS1 and B6 mice

To identify early molecular changes, transcriptional profiles from brain samples from 2-6 months B6.*APP/PS1* and B6 control mice were generated by RNA-seq (Fig. 2A). The initial study cohort included 60 brain samples, and principle components analysis (PCA) identified one brain and one retina sample as outliers which were removed. PCA for the remaining 59 samples is shown in **Fig. S2A**. A generalized linear model (GLM) was used to identify genes that showed expression levels as a function of age, genotype, and age by genotype (Methods; Fig. 2B). All genes were normalized to mean TPM in 2 months B6 samples. First, the expression levels of *APP* and *PSEN1* were assessed. Mapping to the mouse genome was sufficient to detect mutant human *APP* and *PSEN1* transcript expression and therefore, mapping to the human genome was not required. As expected, a significant genotype (APP) effect was observed with both *APP* and *PSEN1* significantly higher in B6.*APP/PS1* compared to B6 brain samples (Fig. 2B). Also, the expression levels of these transgene-driven transcripts did not change significantly across ages assessed (2-6 months). In total, 3969 genes showed significant effects (p < 0.05) in the model (Fig. 3A). Genotype and age by genotype related genes were clustered based on coefficients (*APP*, *APP*4mo, *APP*5mo and *APP*6mo; mo=months). The *APP* group contained 368 genes with increased expression (relative to 2 months B6 samples, herein referred to as ‘upregulated’; **Table S1**) and 317 genes with decreased expression (herein referred to as ‘downregulated’; **Table S2**). (Fig. 3B). Gene set enrichment analysis of the 368 upregulated genes showed overrepresentation (p < 0.1) of multiple Gene Ontology (GO) terms including ‘steroid biosynthesis’, ‘valine, leucine and isoleucine degradation’, ‘gap junction’, ‘butanoate metabolism’ and ‘metabolic pathways’ (Fig. 3C**; Table S3**). The 317 downregulated genes were enriched for ‘protein processing in endoplasmic reticulum’ (**Table S4)**. Enrichment analyses were also performed for genes in the APP4mo (162 upregulated, 164 downregulated), APP5mo (260 upregulated, 325 downregulated) and APP6mo groups (301 upregulated, 315 downregulated) (**Tables S5-16**). ‘Protein processing in endoplasmic reticulum’ was enriched in upregulated genes across all groups (App4mo, APP5mo and APP6mo). In contrast, ‘oxidative phosphorylation’, a process commonly considered disrupted in AD [15], was enriched in downregulated genes across all groups. Transcriptional profiling was performed just prior to or at onset of plaque onset in the B6.*APP/PS1* model, which implies that modulation of metabolic processes, protein processing in the endoplasmic reticulum and oxidative phosphorylation are early events relevant to AD in this model.

**Figure 2.**
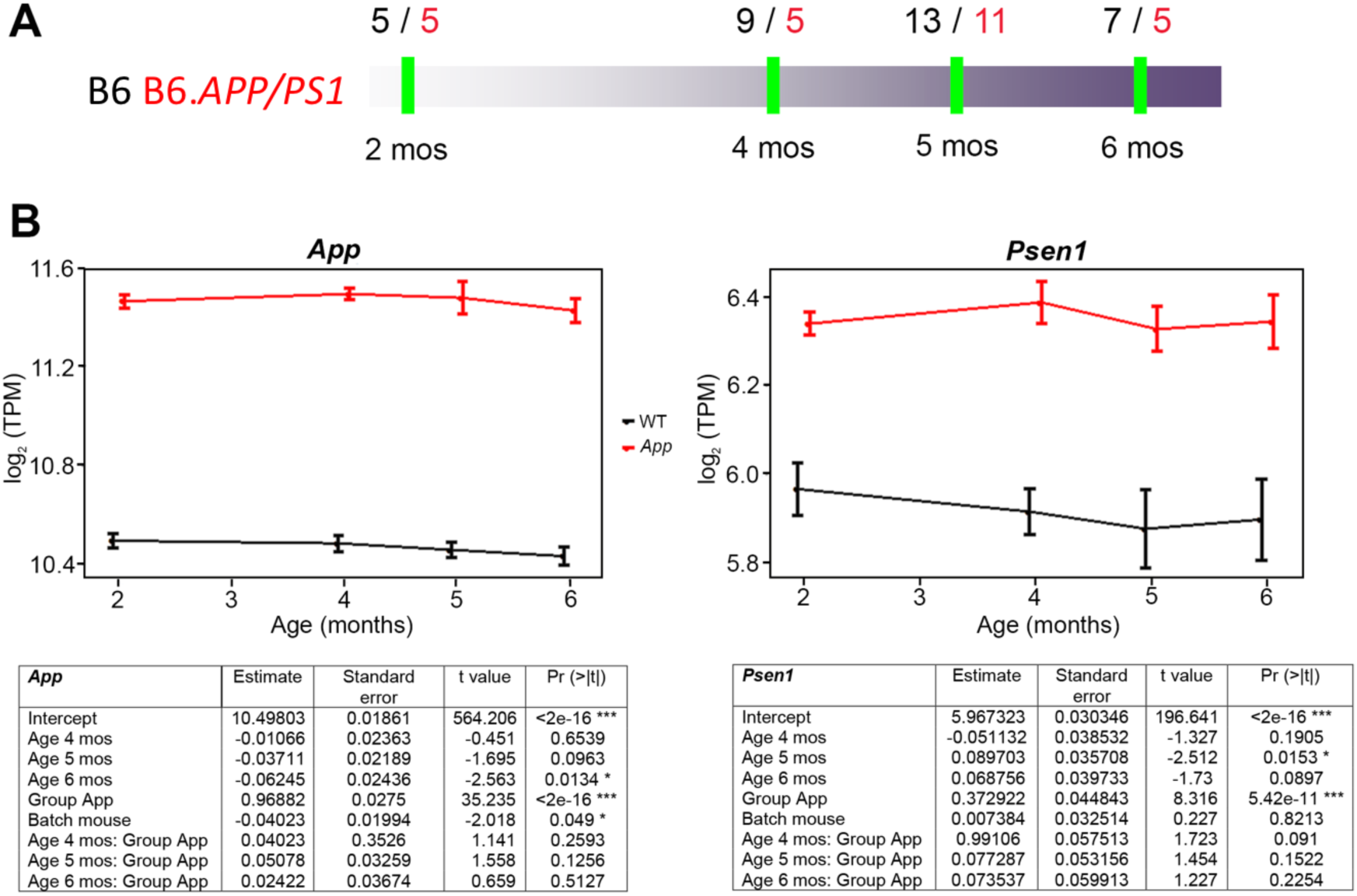
Overview of experimental design. (A) Overview of brain collection for RNA seq analyses. A total of 60 brains were collected at 2, 4, 5 and 6 months age from B6 (WT) and B6.*APP/PS1* female mice. (B) Expression levels of *APP* and *PSEN1* from B6.*APP/PS1* female mouse brains at 2, 4, 5 and 6 months. The longitudinal TPM values estimated from the RNA-seq data are shown. The GLM coefficients for each predictor are also given in tabular format for *APP* and *PSEN1* genes.

**Figure 3.**
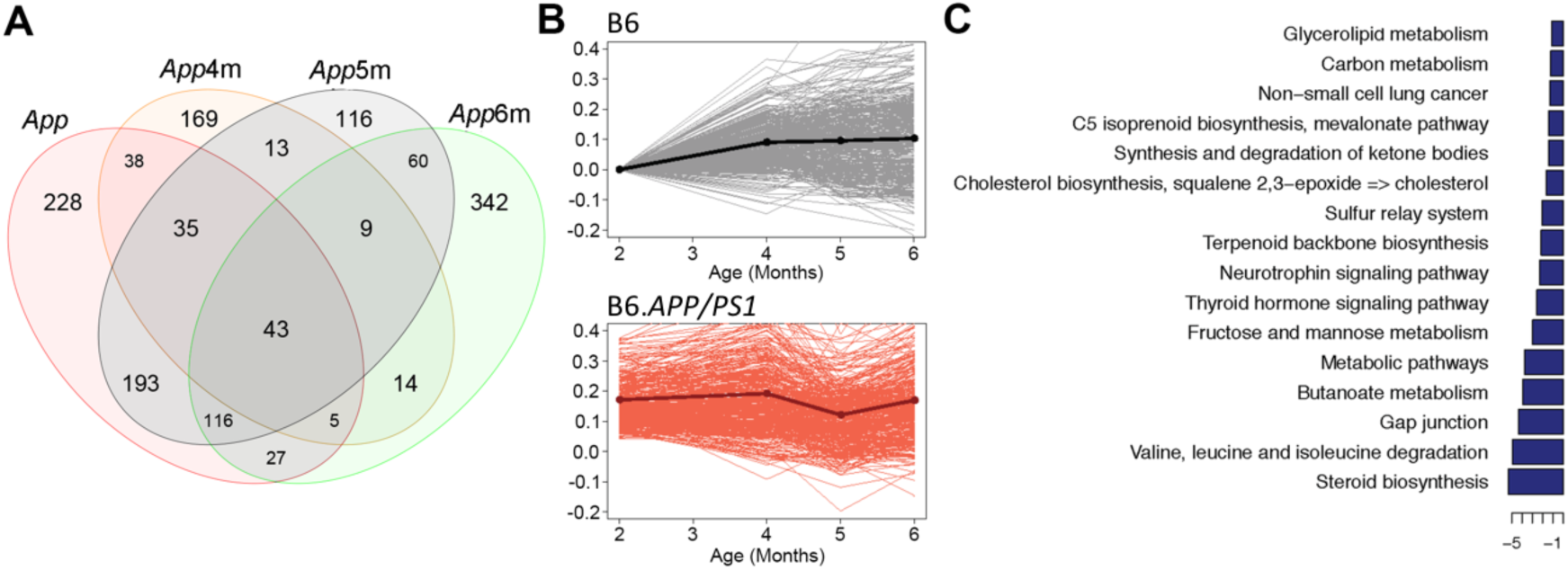
Transcriptomics identifies early changes in B6.*APP/PS1* mouse brains. (A) Venn diagram of numbers of differentially expressed genes (DEG) in B6.*APP/PS1* female mouse brains at 2, 4, 5 and 6 months. (B) Gene clusters based on coefficients from all age groups for B6 (WT) and B6.*APP/PS1* brain. (C) Enriched pathways of genes positively correlated with *APP* from all age groups of brain RNA seq data.

### STAT3 is a master regulator of early responses to plaque deposition in B6.APP/PS1 mice

Previous studies have defined plaque development in B6.*APP/PS1* as a ‘seeding’ phase (prior to 6 months) and an expansion phase (after 6 months) [16, 17]. To understand genes and processes altered at this critical stage of amyloidosis in this model, we further interrogated the APP6mo group. In addition to ‘protein processing in the endoplasmic reticulum’ the 301 upregulated genes in the APP6mo group where enriched for multiple GO terms including ‘ubiquitin mediated proteolysis’, ‘axon guidance’, ‘MAPK signaling’, ‘JAK-STAT signaling pathway’ and ‘endocytosis’ (Fig. 4A-B). The 315 downregulated genes were enriched for GO terms that included ‘oxidative phosphorylation’. ‘non-alcoholic fatty liver disease (NALFD)’ and ‘cardiac muscle contraction’ (**Table S16**). Enrichment of the JAK/STAT pathway was particularly intriguing given multiple studies linking this pathway to amyloidosis and AD (e.g. [18, 19]). Genes in the JAK/STAT pathway included *Crebbp*, *Plxna4*, *Ccdc85b*, *Uba52*, *Rps19*, *Cacna1e*, *Cox17*, *Grid2ip* and *Ntrk3*. Using iRegulon, we predicted Stat3 as a potential master regulator (Fig. 4C). This was consistent with *Stat3* upregulation observed in 6 months B6.*APP/PS1* compared to B6 samples (**Fig. S3**). To identify potential cell types in which JAK/STAT pathway genes were functioning, expression levels of the three genes (*Cox17*, *Ntrk3* and *Grid2ip*) were queried in cell-type specific gene expression data [20]. *Cox17* was expressed in multiple cell types (Fig. 4D) whereas *Ntrk3* was more robustly expressed by astrocytes (Fig. 4E) and Grid2ip, although lowly expressed generally, was biased towards neurons (Fig. 4F). These analyses suggest that the enrichment of JAK/STAT genes may be a result of expression changes in multiple cell types and not simply a single cell type responding to APP/PS1-mediated amyloid deposition.

**Figure 4.**
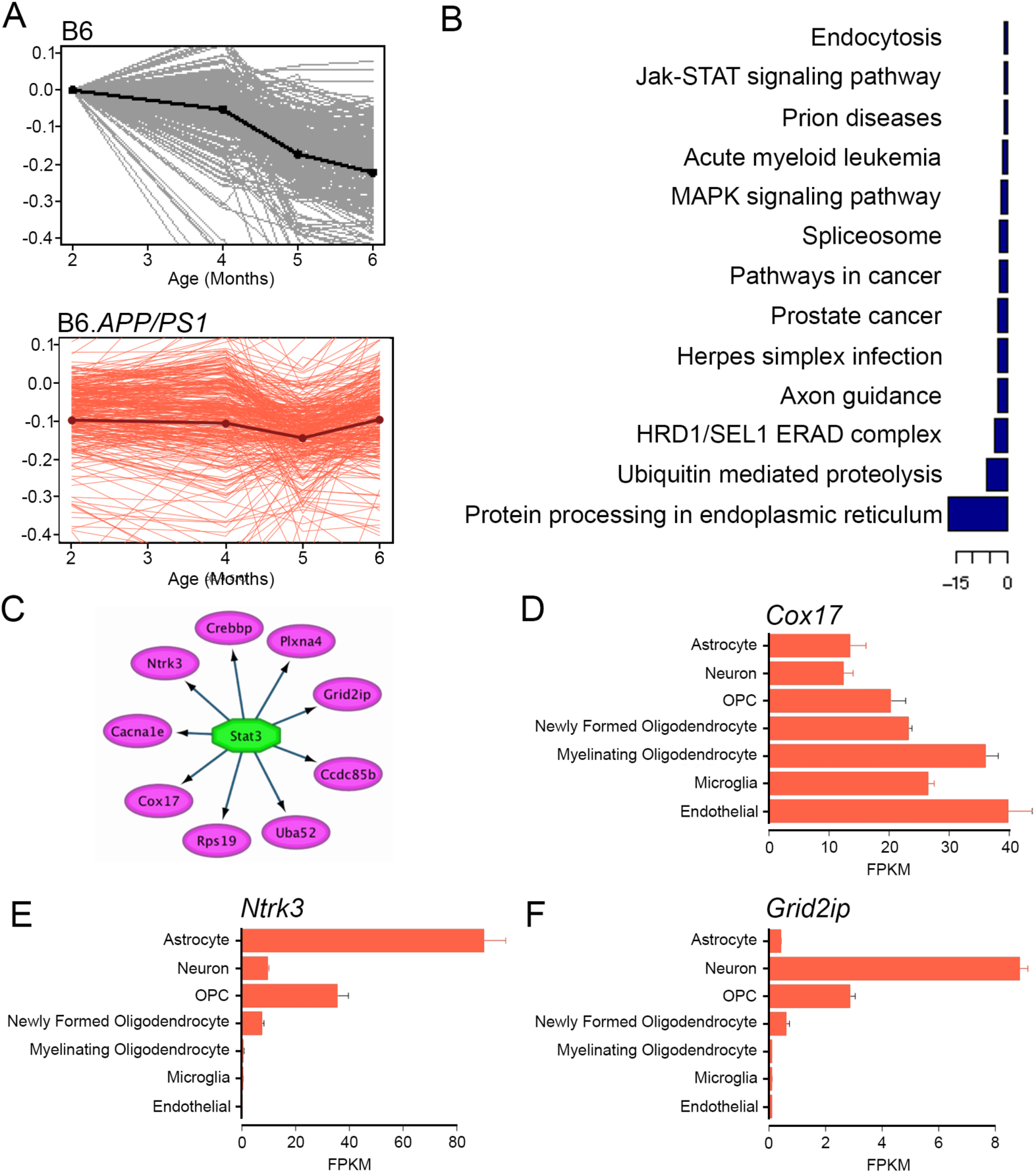
Enrichment of differentially expression genes identifies *Stat3* and associated regulatory network in B6.*APP/PS1* mouse brain. (A) Gene clusters based on coefficients from 6 months old B6 and B6.*APP/PS1* brain. (B) Enriched KEGG pathways of genes at 2, 4, 5 and 6 months positively correlated with *APP* from 6-month group brain RNA seq data. (B) Predicted transcriptional networks for *Stat3* based on iRegulon. Circular nodes represent predicted targets, with *Stat3* as a regulator gene in the center. (D-F) Bar plots of RNAseq data showing FPKM expression of three putative *Stat3* regulated genes (*Cox17*, *Ntrk3*, *Grid2ip*) in astrocyte, neuron, oligodendrocyte progenitor cells (OPC), newly formed oligodendrocytes, myelinating oligodendrocyte, microglia, endothelial cells from P7-P17 mouse brain.

We sought to validate STAT3 as a master regulator of the JAK/STAT genes upregulated in B6.*APP/PS1* compared to B6 mice at 6 months. STAT3 is ordinarily sequestered in the cytoplasm but, upon phosphorylation, is translocated to the nucleus where it can regulate transcription of target genes [21]. Therefore, antibodies to both STAT3 and phosphorylated STAT3 (pSTAT3) were used for localization studies. In brains from 6 months B6.*APP/PS1* mice, STAT3 co-localized with markers of neurons (NEUN), astrocytes (GFAP) and microglia (IBA1) (Fig. 5A). pSTAT3 colocalized with markers of astrocytes and microglia (Fig. 5B). pSTAT3 was not apparent in neurons. Collectively these data suggest a complex role for STAT3 in early stages of disease in B6.*APP/PS1* mice.

**Figure 5.**
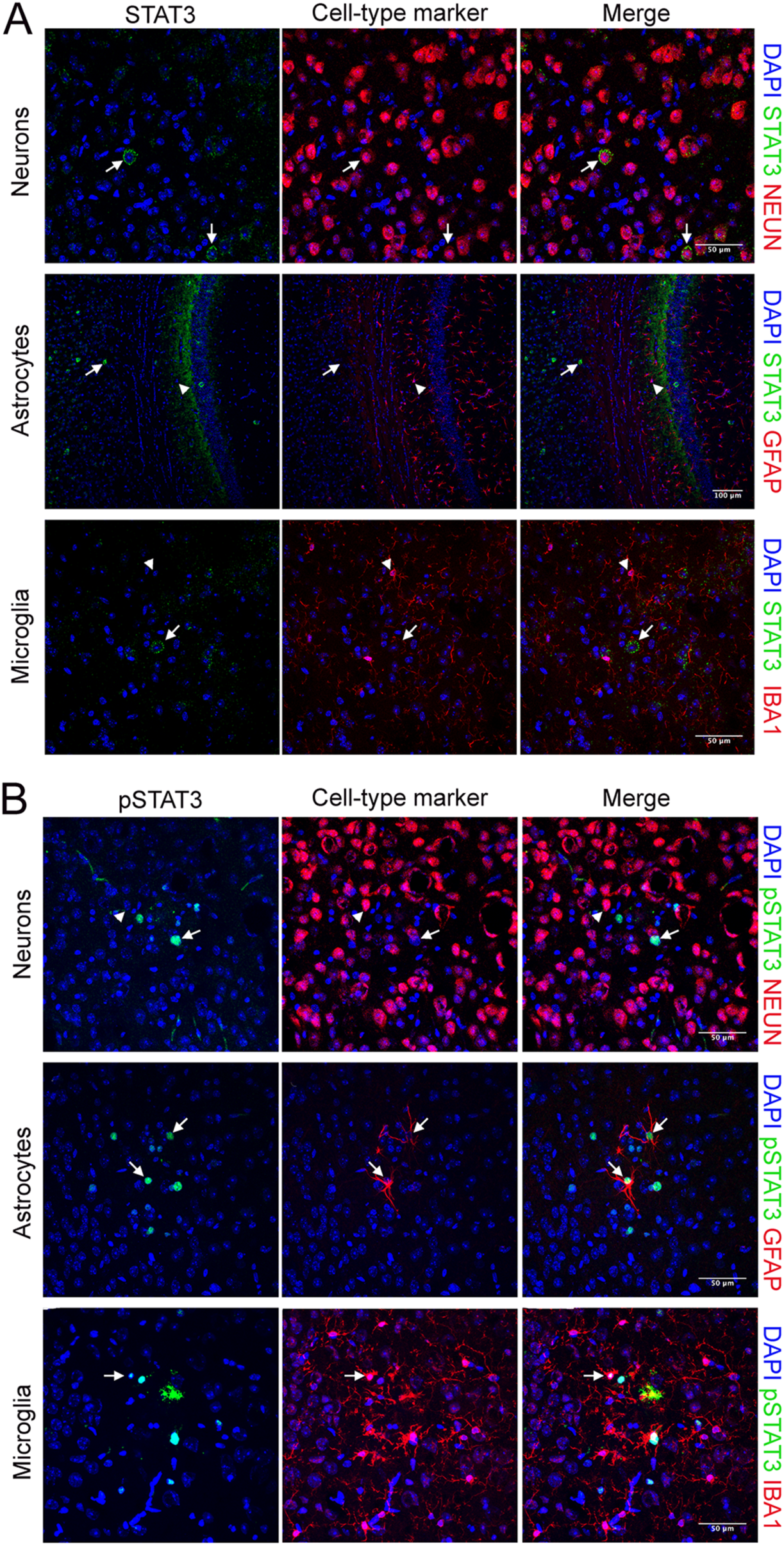
STAT3 is localized to different cell types in B6.*APP/PS1* mouse brain. (A) STAT3 localization was assessed in 6 months B6.*APP/PS1* mouse brain sections. Double labeling of STAT3 (*green*) with cell-type specific markers for astrocytes (GFAP, *red*), microglia (Iba1, *red*), or neurons (NeuN, *red*) in B6.*APP/PS1* mouse brain. STAT3 was localized within cytoplasm of the stained cells. STAT3 was specifically localized in astrocytes in CA1 region. Scale bar: 50 μm, 100 μm. (B) pSTAT3 localization was assessed in 6 months B6.*APP/PS1* mouse brain sections. Double labeling of pSTAT3 (*green*) with cell-type specific markers for astrocytes (GFAP, *red*), microglia (Iba1, *red*), or neurons (NeuN, *red*) in B6.*APP/PS1* mouse brain. pSTAT3 was co-localized with DAPI, suggesting translocation of pSTAT3 to the nucleus. Scale bar: 50 μm.

### Generalized linear modeling of retina samples from B6.APP/PS1 and B6 mice

To determine the potential for the retina to stage changes in the brain, transcriptional profiling of whole retinas from 2 and 6 months B6.*APP/PS1* and B6 control female mice was performed (Fig. 6A). One retina sample had less than 30 million reads and failed basic QC by FastQC. This sample was excluded from further processing, along with another sample PCA identified as an outlier (see Methods). PCA clustering on the remaining 20 samples showed a separation by genotype in retina transcriptome of WT and B6.*APP/PS1* female mice (**Fig. S2B**). GLM analysis was performed using similar criteria to those described for brain samples. Similar to brain samples (Fig. 2B), *APP* and *PSEN1* transcript levels were significantly greater in retinas from B6.*APP/PS1* compared to B6 controls (**Fig. S4)**. This indicates that the *APP/PS1* transgenes are active in the retina and the mutant transcript levels do not change significantly at the ages assessed. A total of 9021 genes were identified using GLM. A total of 2596 genes were identified in the APP group (differentially expressed in B6.*APP/PS1* compared to B6, independent of age) and 466 identified in the APP6mo group (Fig. 6B). All but 2 of the 466 genes in the APP6mo group were also present in the APP group indicating that the vast majority of transcriptional changes (2130 genes) in the retina occur at or prior to 2 months. Gene set enrichment was performed on the 850 upregulated (**Tables S17, S19**) and the 1280 downregulated (**Tables S18, S20**) genes in the APP group. The upregulated genes were enriched for processes seen in the brain (e.g. ‘oxidative phosphorylation’, ‘NAFLD’, ‘metabolic pathways’ and cardiac muscle contraction) (Fig. 6C-D) although these were downregulated in the brain samples. Analyses of the 1280 downregulated genes identified enriched GO terms related to synapse biology (‘GABAergic synapse’, ‘dopaminergic synapse’, ‘glutamatergic synapse’ and ‘synaptic vesicle cycle’).

**Figure 6.**
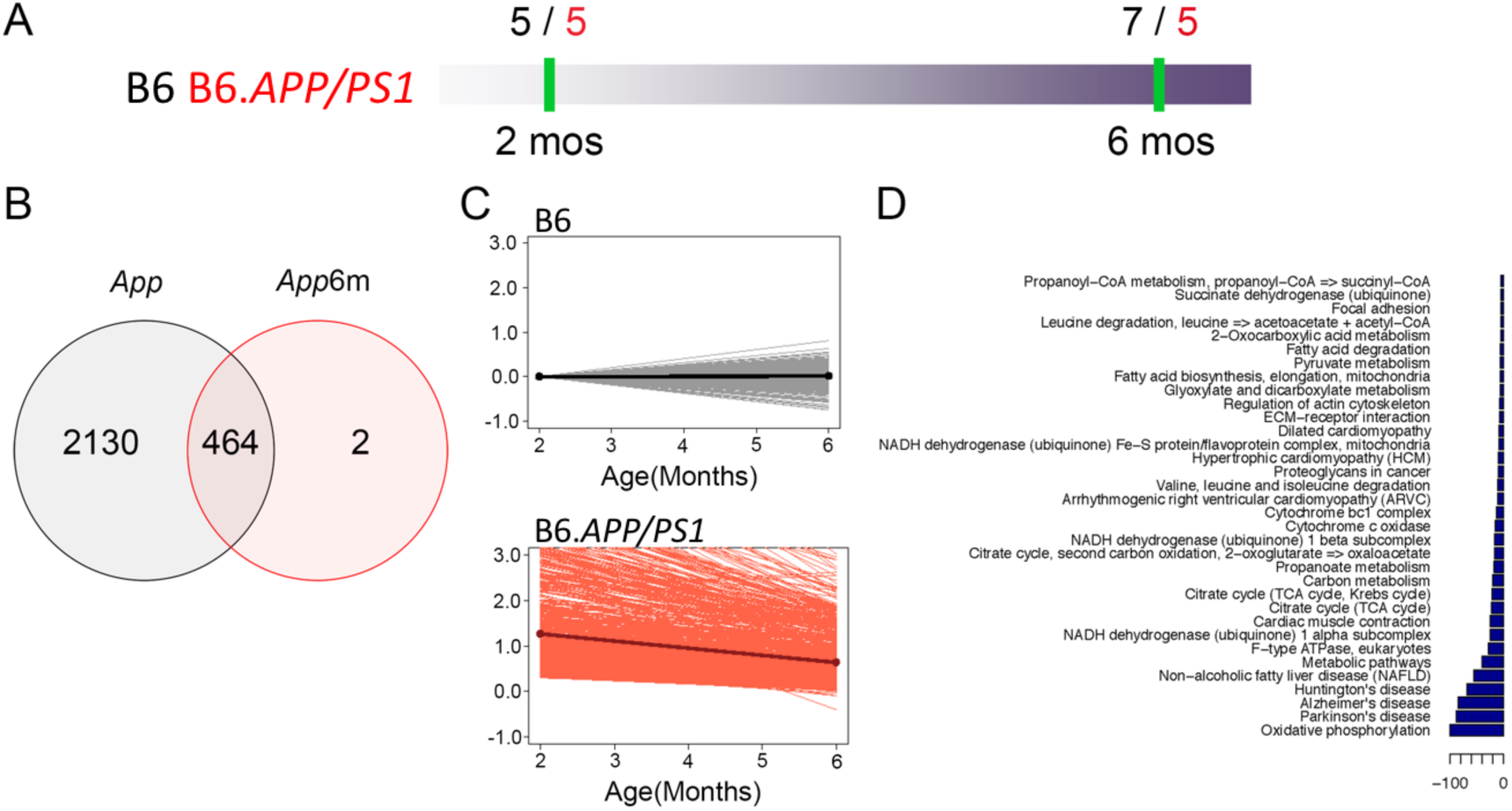
Transcriptomics identifies early changes in B6.*APP/PS1* mouse retinas. (A) Overview of retina collection for RNA seq analyses. A total of 22 retinas each collected at 2 and 6 months B6 (WT) and B6.*APP/PS1* female mice. (B) Venn diagram of numbers of differentially expressed genes (DEG) in B6.*APP/PS1* female mouse retinas at 2 and 6 months. (C) Gene clusters based on coefficients from 2 and 6 months groups for B6 (WT) and B6.*APP/PS1* retina. (D) Enriched pathways of genes positively correlated with APP/PS1 (*APP*) from 6 months group of retina RNA seq data.

Only 151 genes were upregulated in the APP6mo group (**Table S21, S23**). However, 315 genes (**Table S22**) were downregulated and enriched for ‘dilated cardiomyopathy’, ‘hypertrophic myopathy’, ‘fatty acid degeneration’ and ‘tight junction’ (**Fig. S5 Table S24**). Additional vascular-related GO terms including ‘adrenergic signaling on cardiomyocytes’, ‘arryhthmogenic right ventricular cardiomyopathy’ (ARVC), ‘ECM-receptor interaction’ and ‘focal adhesion’ were also enriched, suggesting vascular perturbations in the retinas of B6.*APP/PS1* mice. For further analysis, retinal cross sections from B6.*APP/PS1* and B6 were stain with fibrinogen and CD31 antibodies. Microhemorrhages were observed in large and small blood vessels within the retinas of B6.*APP/PS1* but not B6 controls (Fig. 7). Interestingly, this appeared somewhat age-dependent with retinas of 6 months B6.*APP/PS1* mice appearing to show more retinal microhemorrhages than those from 2 and 4 months B6.*APP/PS1* mice (Fig. 7).

**Figure 7.**
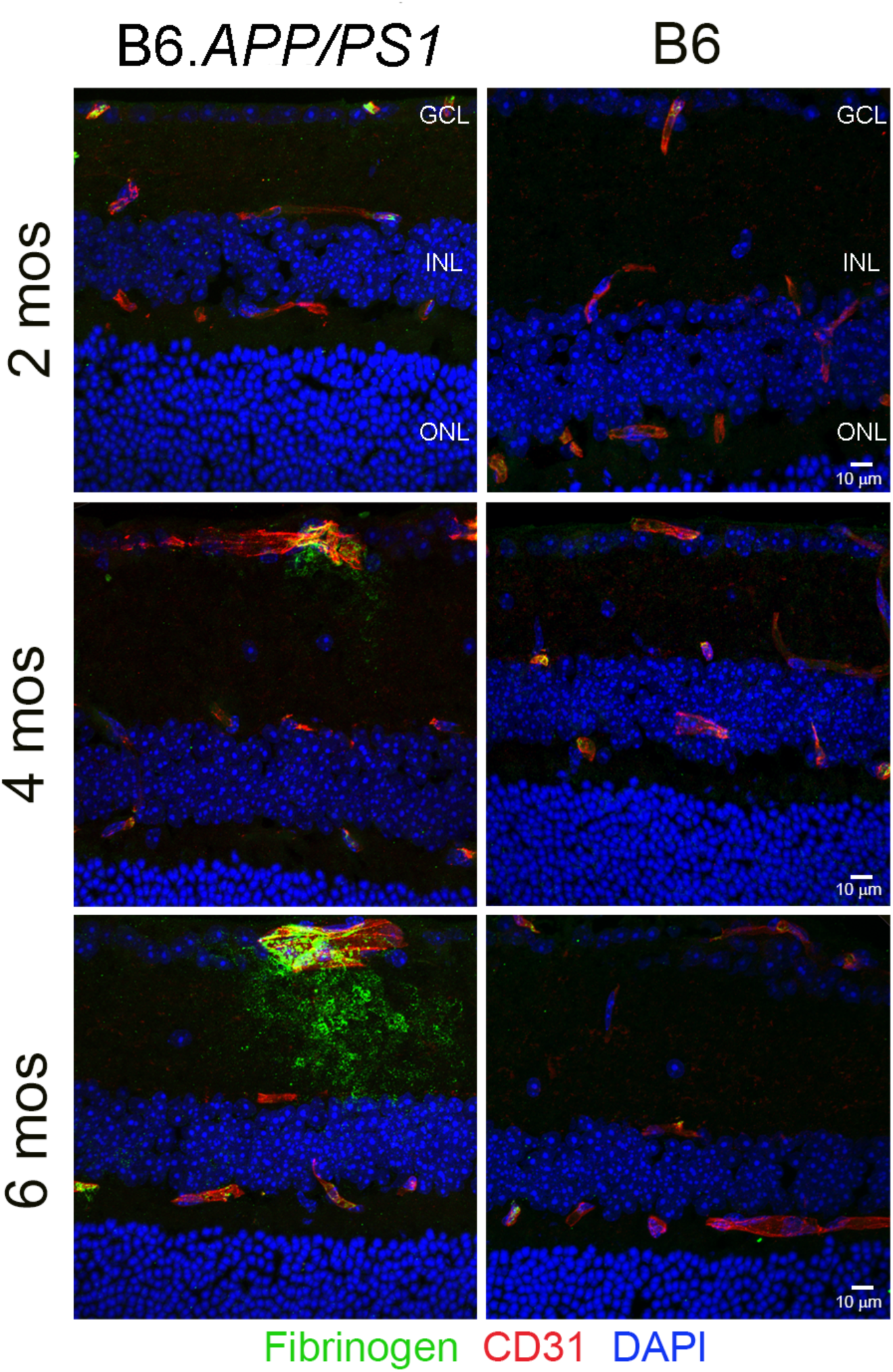
Evidence of microvascular microhemorrhages in B6.*APP/PS1* mouse retina. Vascular leakage in 2, 4 and 6 months old B6.*APP/PS1* and B6 control retinas were assessed via detection of fibrin(ogen) in the retina. Confocal images revealed fibrin(ogen) (*green*) immunoreactive areas near blood vessels stained with CD31 (*red*) in retinal cross sections at 4 and 6 months old B6.*APP/PS1* mice, but not in age matched controls. In most blood vessels, fibrin(ogen) staining was localized to within the lumen of blood vessels. However, in some major blood vessels, fibrin(ogen) staining extended beyond the vascular margin, creating a fuzzy leakage pattern, indicative of loss of vascular integrity. Fibrinogen leakage was noticeably worse at 4 and 6 months compared to 2 months old B6.*APP/PS1* mice and B6. Abbreviations: GCL, ganglion cell layer; INL, inner nuclear layer; ONL, outer nuclear layer. n=4-6; Scale bar: 10 μm

## DISCUSSION

Here we used a generalized linear model (GLM) to identify sets of genes that show differential expression patterns just prior to and during the onset of plaque deposition in B6.*APP/PS1* mice, a commonly used mouse model relevant to AD. Brain samples were assessed from 2 months in both B6.*APP/PS1* and B6 controls, which allowed for the identification of genes that are differentially expressed prior to plaque onset. Our approach of transcriptional profiling at multiple ages around plaque onset allowed us to identify changes that were due to age, genotype and age-by-genotype. However, the challenge in using mouse models based on familial mutations is to identify molecular changes that are relevant to early stages of human AD. Any treatment for AD is likely to be most effective when delivered as early in the disease progression as possible. In B6.*APP/PS1*, amyloid deposition is driven by two transgenes that generate high levels of both mutant APP (APP^swe^) and PSEN1 (PSEN1^de9^). This has been shown to cause phenotypes that are considered to be artefactual and likely not relevant to human AD. For instance, B6.APP/PS1 mice show hyperactivity and increased seizures compared to their B6 counterparts – attributed to high levels of expression of the APP gene. Expressing the transgenes later using conditional expression strategies eliminates these possible side effects [22], although is also unlikely to accurately recapitulate the pathogenesis of late-onset AD. Therefore, caution should be taken when using these mice for identifying genes/pathways that precede plaque onset.

The experimental design allowed the identification of groups of genes that were changing across the ages when plaques were seeding. This timeframe has been previously defined in B6.*APP/PS1* as between 4-6 months. Therefore, we were particularly interested in genes that were differentially expressed at 6 months. These sets of genes are potentially most relevant to early stages of plaque deposition in humans. Gene set enrichment identified multiple biological processes that are impacted by overexpression of *APP/PS1* in this age range. Oxidative phosphorylation and metabolic pathways appear disrupted at this early stage and have been associated with early stages of human AD [23, 24]. In addition, non-alcoholic liver fatty liver disease (NAFLD) was also identified at this early stage and has been previously associated with AD [25, 26]. NAFLD is generally characterized by excessive fat build up in the liver. White matter tracts have a very high lipid to protein ratio (lipids account for at least 70% of dry weight) and therefore white matter regions require high levels lipids, such as cholesterol. The identification of the NAFLD pathway might suggest previously unidentified white matter changes early in B6.*APP/PS1* mice. The potential for white matter changes making a greater contribution to the early pathophysiology of Alzheimer’s disease are now being explored [27]. Intriguingly, brain myelination transcriptional networks were down regulated in AD, although more strikingly in progressive supranuclear palsy (PSP) [28]. Therefore, a previously underappreciated role of oligodendrocytes requires further investigation. Environmental risk factors for AD such as a western diet and obesity also cause inflammation and changes to white matter damage and changes to oligodendrocytes (e.g. [29–31]). Therefore, given the current obesity epidemic, treatments that target white matter changes and oligodendrocyte function are potentially crucial to combat AD and related dementias.

A pathway that this study and others [19, 32] have identified as potentially a modulator of plaque deposition is the JAK/STAT (Janus kinas/signaling transducers and activators of transcription) pathway. The JAK/STAT pathway is involved in immunity, cell division and death, tumor formation and multiple diseases/injuries. Central to the JAK/STAT pathway are the STAT genes. STATs are transcription factors that are commonly activated to regulate gene transcription in response to cytokines and growth factors. A recent study showed the importance of STAT3 in plaque clearance by microglia and identified interleukin-10 as an upstream signaling molecule [19, 32]. However, STAT3 and pSTAT3 are expressed by multiple cell types including neurons, astrocytes and microglia. Glial cells in particular are likely to play both beneficial and damaging roles at different stages of AD. For instance, although microglia may be required to phagocytose and/or clear amyloid, astrocytes may provide a protective barrier to prevent the spread of toxic molecules. In other contexts, such as spinal cord injury, inhibiting STAT3 in astrocytes was actually damaging [33]. Therefore, identifying cell-specific downstream targets of STAT3 may be a viable strategy to identify more precise treatments. The analysis performed in this study identified nine potential downstream targets of STAT3 including *Cox17*, *Ntrk3* and *Grid2ip*. According to cell-specific gene expression resources, *Ntrk3* is more highly expressed in astrocytes and may provide a target to test the astrocyte-specific role of STAT phosphorylation in amyloid deposition and clearance.

Much attention is now being given to using the eye, particularly the retina, as a window to the brain (e.g. [34–37]). Given the similarity in retinal and brain tissue there is great potential for staging early AD-relevant changes in the retina. This has the advantage that, in contrast to brain imaging, the retina can be readily imaged for little cost multiple times throughout a person’s lifetime. Those considered to be at risk based on changes in the retina could be prioritized for more in-depth studies such as PET/MRI and/or blood or CSF biomarkers. However, retinal imaging for AD is controversial. While some studies in both humans and mouse models show AD-relevant changes in the retina including amyloid and tau deposition, retinal ganglion cell dendrite changes [34, 38–40], others report little to no changes in the retina [6]. Here, we assessed retinal changes by both histology and transcriptional profiling. Amyloid deposition was observed but appeared quite different to the amyloid in the brain. Retinal amyloid occurred more often closely associated with vessels whereas vascular amyloid is rare in the brain in this model – at least on a B6 background [41]. This may suggest differences in the ability of retinal and brain cells to process amyloid. Some reports suggest retinal neurons express higher levels of alpha-secretase which may allow greater processing of mutant forms of APP to a non-amyloidogenic form. Interestingly, soluble APP, a non-amyloidogenic processing product, has been shown to be neuroprotective in retinal rotenone toxicity [42]. Transcriptional profiling identified differential expression of genes associated with vascular health in B6.*APP/PS1* compared to B6 samples. Vascular leakage was confirmed using immunofluorescence. Vascular changes were not predicted by transcriptional profiling in the brain and have not been reported in this model. However, the majority of genes were differentially expressed in B6.*APP/PS1* compared to B6 samples at both 2 months and 6 months of age. This may indicate retinal changes due to the overexpression of *APP/PS1* may be due to perturbations to later stages of retinal development/maturation as opposed to early stages of amyloid deposition that would be useful to stage early changes in the brain.

A major limitation of the B6.*APP/PS1* when considering identifying genes/pathways relevant to early stages of plaque development is that plaques begin to develop in young mice. This is motivating multiple groups to develop new models that develop plaque deposition in a more age-dependent manner. Approaches include using targeted replacement of the mouse *App* gene with both the human *APP* gene as well as mutant forms of human *APP* [43]. Assessing mice carrying human *MAPT* is also underway. In addition, combining multiple risk factors for sporadic or late-onset AD (such as *APOE^E4^* and *TREM2^R47H^*) is also being assessed. Ultimately, if these new models develop age-dependent AD phenotypes such as amyloid deposition, tau accumulation, neuroinflammation and neuronal cell loss, findings in these next generation of mouse models of AD should provide improved translatability to human AD due their improved construct validity. These would then be the ideal models to assess the full potential for using the eye as a biomarker for tracking early stages of AD and related dementias.

## ACKNOWLEDGEMENTS

This study is part of the Model Organism Development and Evaluation for Late-onset Alzheimer’s Disease (MODEL-AD) consortium funded by the National Institute on Aging. MODEL-AD comprises the Indiana University/The Jackson Laboratory MODEL-AD Center U54 AG054345 led by Bruce T. Lamb, Gregory W. Carter, Gareth R. Howell, and Paul R. Territo and the University of California, Irvine MODEL-AD Center U54 AG054349 led by Frank M. LaFerla and Andrea J. Tenner. This work was also supported by The Pyewacket Foundation (G.W.C.); and The Jackson Laboratory Director’s Innovation Fund (G.W.C. and G.R.H.). The funding organizations played no role in the design and conduct of the study; in the management, analysis, and interpretation of the data; or in the preparation, review, or approval of the article.

## Authors’ contributions

G.R.H and G.W.C were responsible for the study concept and design. S.R.C, A.U, H.W.J and C.A were responsible for data collection and executed experiments. G.R.H, G.W.C, S.R.C, X.W. and A.U analyzed and interpreted the data. S.R.C, G.R.H, G.W.C, and A.U drafted the manuscript. G.R.H, G.W.C, S.R.C, A.U, M.S, H.W.J and C.A revised the manuscript. G.R.H and G.W.C obtained the funding and were responsible for the study supervision.

## DISCLOSURE STATEMENT

The authors have no conflict of interest to report.

