## Supplemental figures for "Staging Alzheimer’s disease in the brain and retina of *B6.APP/PS1* mice by transcriptional profiling"

### SUPPLEMENTAL MATERIAL

### SUPPLEMENTAL FIGURES

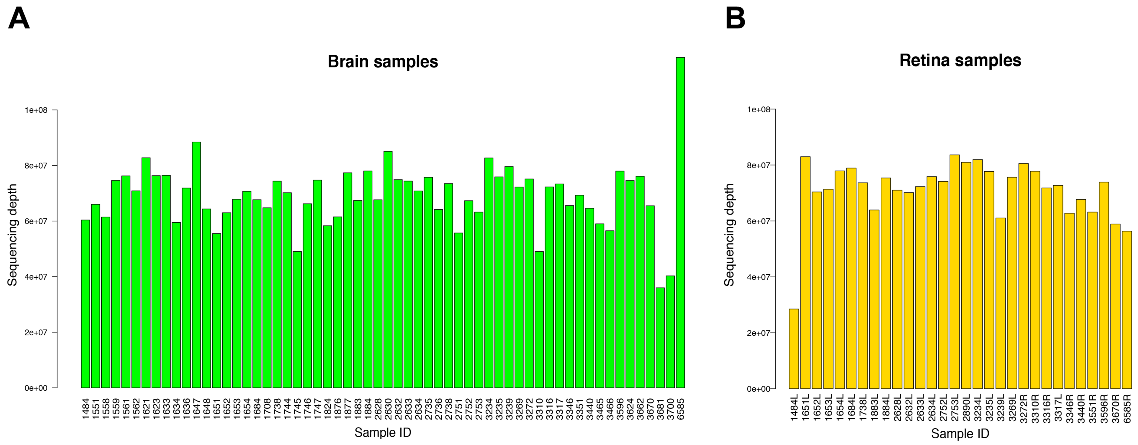


**Supplementary Figure 1**

Sequencing depths in terms of number of short sequence reads as calculated from fastq files for brain (A) and retina (B) samples.

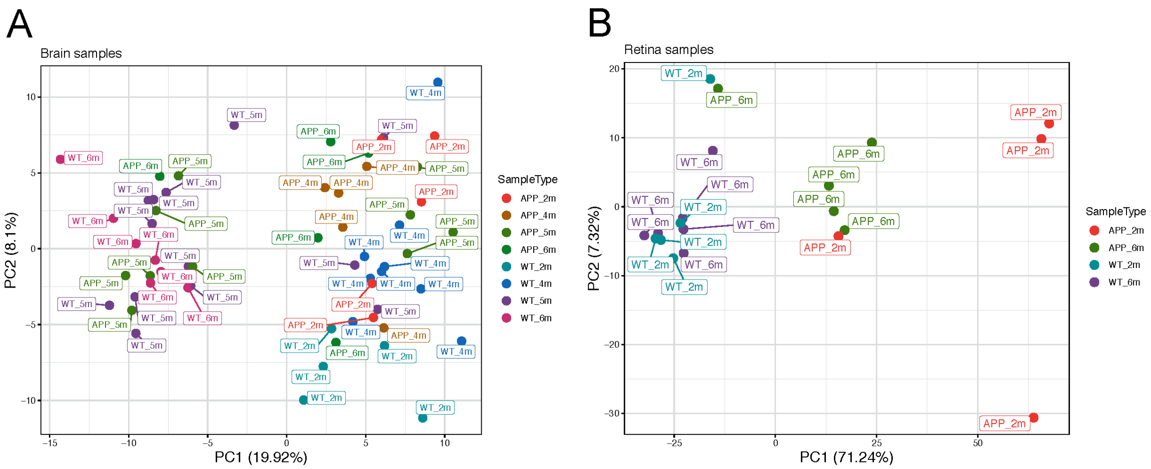


**Supplementary Figure 2**

Principal Component Analysis (PCA) clustering of RNA-Seq transcriptomes from brain (A) and retine (B) samples.

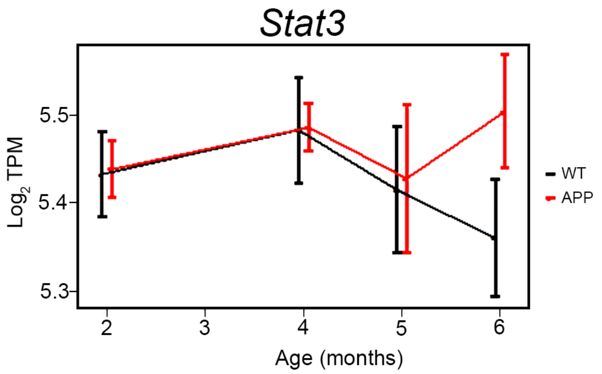


| *Stat3* | Estimate | Standard error | t value | Pr (>\|t\|) |
|---|---|---|---|---|
| Intercept | 5.431993 | 0.029024 | 187.154 | <2e-16 *** |
| Age 4 mos | 0.050087 | 0.036853 | 1.359 | 0.1802 |
| Age 5 mos | -0.017509 | 0.034153 | -0.513 | 0.6104 |
| Age 6 mos | -0.072281 | 0.038002 | -1.902 | 0.0629 |
| Group App | 0.005949 | 0.04289 | 0.139 | 0.8902 |
| Batch mouse | -0.058435 | 0.031097 | -1.879 | 0.0661 |
| Age 4 mos: Group App | -0.002873 | 0.055007 | -0.052 | 0.9586 |
| Age 5 mos: Group App | 0.006968 | 0.05084 | 0.137 | 0.8915 |
| Age 6 mos: Group App | 0.137732 | 0.057303 | 2.404 | 0.02 * |

**Supplementary Figure 3**

Expression level of *Stat3* from B6.*APP/PS1* female mouse retina at 2 and 6 months. The longitudinal TPM values estimated from the RNA-seq data are shown. The GLM coefficients for each predictor are also given in tabular format.

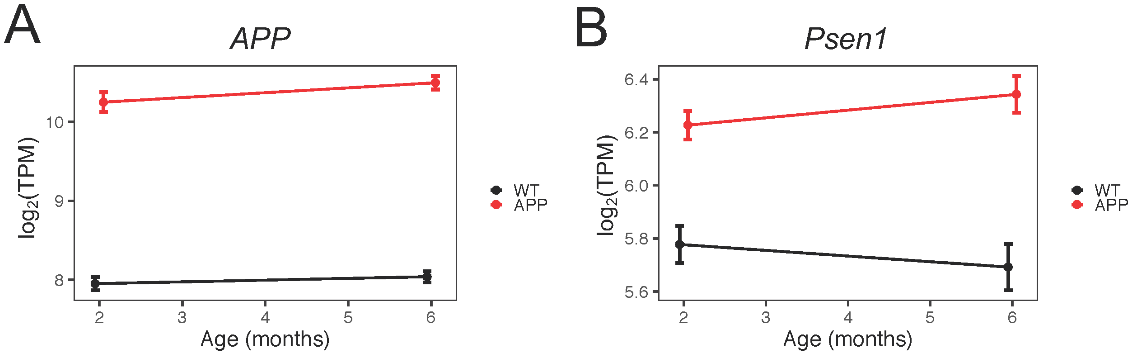


| ***APP*** | Estimate | Std. Error | t value | Pr(>\|t\|) |
|---|---|---|---|---|
| (Intercept) | 7.94953 | 0.04187 | 189.868 | < 2e-16 |
| age6m | 0.08831 | 0.05669 | 1.558 | 0.1401 |
| groupAPP | 2.30191 | 0.07184 | 32.04 | 3.14e-15 |
| batchmouse | -0.09103 | 0.06978 | -1.304 | 0.2117 |
| age6m:groupAPP | 0.15792 | 0.08716 | 1.812 | 0.0901 |

| ***Psen1*** | Estimate | Std. Error | t value | Pr(>\|t\|) |
|---|---|---|---|---|
| (Intercept) | 5.77747 | 0.03383 | 170.774 | < 2e-16 |
| age6m | -0.085333 | 0.04581 | -1.863 | 0.0822 |
| groupAPP | 0.44956 | 0.05805 | 7.744 | 1.28e-06 |
| batchmouse | -0.05475 | 0.05639 | -0.971 | 0.3469 |
| age6m:groupAPP | 0.20139 | 0.07042 | 2.860 | 0.0119 |

**Supplementary Figure 4**

Expression levels of *APP* and *PSEN1* from B6.*APP/PS1* female mouse retina at 2 and 6 months. The longitudinal TPM values estimated from the RNA-seq data are shown. The GLM coefficients for each predictor are also given in tabular format for *APP* and *PSEN1* genes.

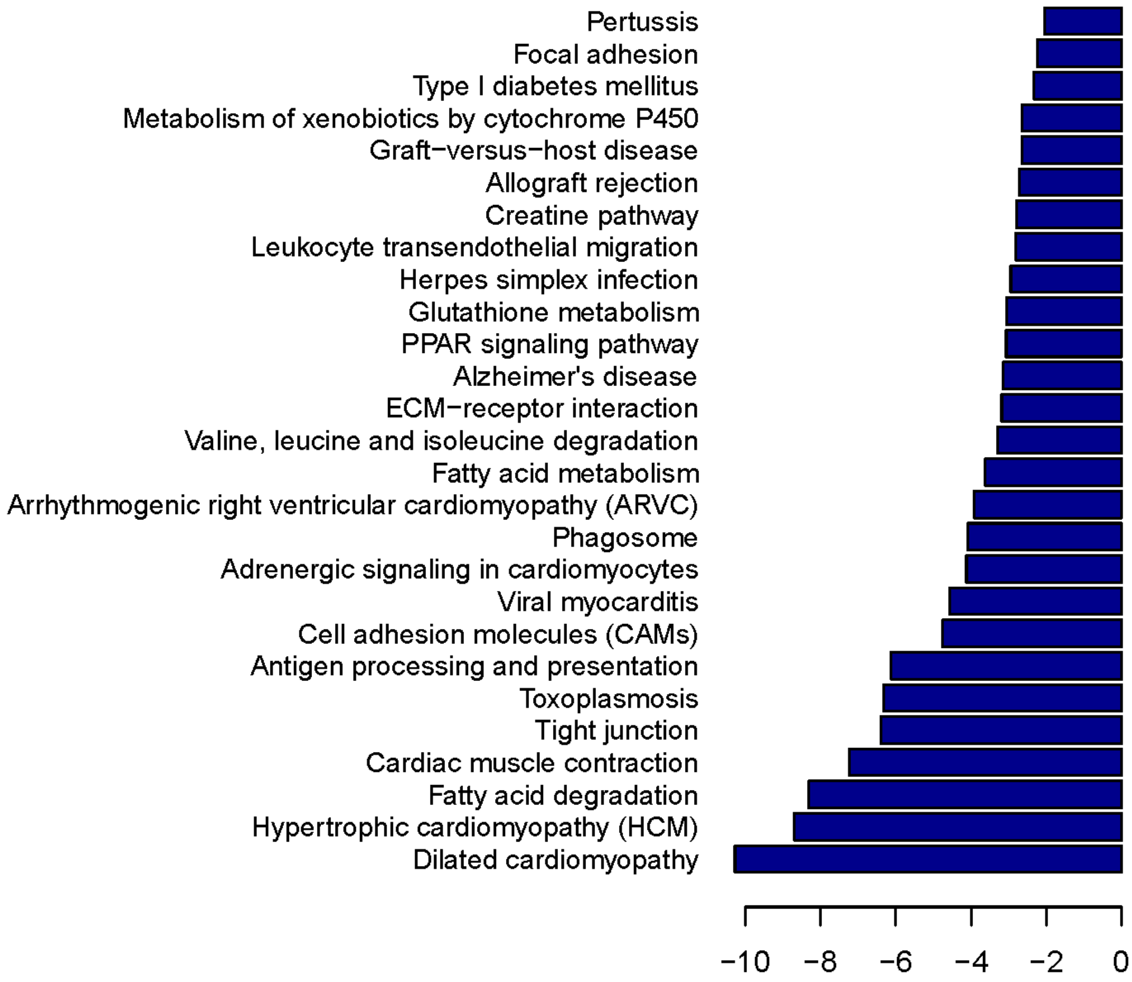


**Supplementary Figure 5**

Enriched pathways of genes negatively correlated with *APP* at 6 months of age in retina RNA seq data.

### SUPPLEMENTAL TABLES

**Brain datasets:**
**APP group:**
**Table S1:** Upregulated genes in APP group
**Table S2:** Downregulated genes in APP group
**Table S3:** Gene set enrichment analysis of upregulated genes in APP group
**Table S4:** Gene set enrichment analysis of downregulated genes in APP group

**APP4mo**
**Table S5:** Upregulated genes in APP4mo
**Table S6:** Downregulated genes in APP4mo
**Table S7:** Gene set enrichment analysis of upregulated genes in APP4mo
**Table S8:** Gene set enrichment analysis of downregulated genes in APP4mo

**APP5mo**
**Table S9:** Upregulated genes in APP5mo
**Table S10:** Downregulated genes in APP5mo
**Table S11:** Gene set enrichment analysis of upregulated genes in APP5mo
**Table S12:** Gene set enrichment analysis of downregulated genes in APP5mo

**APP6mo**
**Table S13:** Upregulated genes in APP6mo
**Table S14:** Downregulated genes in APP6mo
**Table S15:** Gene set enrichment analysis of upregulated genes in APP6mo
**Table S16:** Gene set enrichment analysis of downregulated genes in APP6mo

**Retina datasets:**
**APP group:**
**Table S17:** Upregulated genes in APP group
**Table S18:** Downregulated genes in APP group
**Table S19:** Gene set enrichment analysis of upregulated genes in APP group
**Table S20:** Gene set enrichment analysis of downregulated genes in APP group

**APP6mo**
**Table S21:** Upregulated genes in APP6mo
**Table S22:** Downregulated genes in APP6mo
**Table S23:** Gene set enrichment analysis of upregulated genes in APP6mo
**Table S24:** Gene set enrichment analysis of downregulated genes in APP6mo
